# ER-Localized Ceramide Accumulation Contributes to Replicative Senescence

**DOI:** 10.1101/2025.05.16.654528

**Authors:** Shweta Chitkara, Mengru Li, Natasha Gozali, Andrey Kuzmin, Artem Pliss, Paras Prasad, Yasemin Sancak, G. Ekin Atilla-Gokcumen

**Author notes:** Corresponding authors (G. Ekin Atilla-Gokcumen), (Yasemin Sancak).

## Abstract

Ceramides regulate diverse cellular processes through compartment-specific accumulation. While mitochondrial ceramide accumulation promotes apoptosis, its regulation and function during senescence remain incompletely understood. Here, we integrate lipidomics, transcriptomics, Raman spectroscopy and biochemical characterizations to define sphingolipid remodeling in replicative senescence. Senescent cells exhibit elevated ceramide levels and depletion of very-long-chain sphingomyelins, despite unaltered sphingomyelin synthase 1 expression, implicating impaired ceramide-sphingomyelin turnover. Pharmacological inhibition of ceramide transfer protein (CERT), the ER-to-Golgi ceramide transporter, phenocopies this sphingolipid remodeling and enhances senescence markers, suggesting disrupted ceramide trafficking as a driver of senescence. Raman spectroscopy suggests ceramide accumulation localized to the ER, with no enrichment at mitochondria, consistent with a non-apoptotic phenotype. Analysis of ER-enriched fractions confirm increased ceramide levels in ER fractions of senescent cells. Mechanistically, ceramide accumulation at the ER can contribute to ER stress, a key component of the senescence program. These findings identify altered ceramide trafficking as a contributor to ER stress and highlight ER-localized ceramide as a critical component of senescence-associated sphingolipid remodeling.

**Significance:** How ceramides function during replicative senescence has remained poorly understood. In this study, we demonstrate that senescent cells accumulate ceramides at the endoplasmic reticulum (ER), revealing a fundamentally different mode of ceramide regulation and function in senescence. By integrating lipidomics, enzymatic assays, Raman-BCA imaging, and biochemical assays, we show that impaired ER-to-Golgi ceramide transport leads to ER-localized ceramide retention. Disrupting ceramide trafficking through inhibition of the ceramide transfer protein CERT phenocopies senescence-associated lipid remodeling and enhances senescence markers, establishing altered ceramide transport as a mechanistic contributor to this process. These findings link impaired ceramide trafficking to replicative senescence and suggest that ceramide function in senescence is linked to its retention at the ER.

## INTRODUCTION

Ceramides are central molecules in sphingolipid metabolism and are synthesized *de novo* through a highly regulated enzymatic pathway^1, 2^. The process begins at the endoplasmic reticulum (ER) membrane with the condensation of serine and palmitoyl-CoA, catalyzed by serine palmitoyltransferase (SPT), producing 3-ketosphinganine (**Scheme 1A**). This intermediate is reduced to sphinganine and subsequently *N*-acylated with different fatty acyl chains by ceramide synthases (CerSs), generating dihydroceramides. These dihydroceramides are then desaturated by dihydroceramide desaturases (DESs) to form ceramides. In addition to *de novo* synthesis, ceramides can also be generated via the hydrolysis of sphingomyelin by sphingomyelinases (SMases) or through the salvage pathway, where sphingosine, derived from the breakdown of complex sphingolipids, is re-acylated by CerS. Once formed, ceramides serve as precursors for complex sphingolipids or other bioactive lipids. Tightly controlled synthesis and metabolism of ceramides within distinct subcellular compartments ensure their availability and functional specificity (reviewed in^2^).

Ceramides have attracted significant attention as they are involved in diverse cellular processes^3–6^. The mechanisms and consequences of ceramide accumulation have been most extensively studied in the context of apoptosis. Elevated ceramide levels at the mitochondria have been shown to disrupt mitochondrial integrity, permeability, and function, thereby promoting cell death^7^. Notably, ceramides localized to the mitochondria induces apoptosis, emphasizing that the cellular localization of ceramides can be linked to their distinct roles in various cellular processes^8^. Mechanistically, ceramides can promote apoptosis by forming ceramide channels in the outer mitochondrial membrane, which facilitate the release of cytochrome c and other pro-apoptotic factors^7,9^. They also disrupt mitochondrial membrane integrity by interacting with key regulatory enzymes of the Bcl-2 protein family^10^. For example, ceramides inhibit phosphoinositide-3-kinase and Akt/PKB signaling, leading to the dephosphorylation and activation of pro-apoptotic proteins such as BAD^11^. Furthermore, ceramides can facilitate the translocation and oligomerization of BAX at the mitochondria, compromising membrane integrity, and interacting with voltage-dependent anion channels, amplifying apoptotic signaling^12^. These studies collectively underscore the critical role of mitochondrial ceramide accumulation in mediating specific interactions at the mitochondria that ultimately drive apoptosis (reviewed in^2^).

In addition to apoptosis, ceramides accumulate in replicative senescence, a state of irreversible cell cycle arrest. Senescence is driven by stress-induced pathways, including the p53-p21 and p16-RB axes, which inhibit cyclin-dependent kinases, preventing cell cycle progression. Several studies have reported ceramide accumulation during replicative^13, 14^ and other senescence programming^15, 16^, implicating these lipids in aging and age-related diseases^17^. Studies have demonstrated that genes involved in sphingolipid metabolism are enriched among transcripts that are differentially expressed during senescence^13, 14^, ceramides accumulate with aging^4, 18^, can induce senescence^18, 19^, and inhibition of *de novo* ceramide biosynthesis reduced senescence^20^. Overall, these studies suggest a critical role for ceramides in senescence. However, the mechanisms by which ceramides and sphingolipids are regulated and function in senescence remain poorly understood.

In this study, we investigate the regulation of ceramides and other sphingolipids during replicative senescence compared to apoptosis (**Scheme 1B**). Through comparative lipid profiling and gene expression analysis, we identify potential distinct sphingolipid regulatory patterns and candidate enzymes that modulate ceramide levels in senescence. The lipidomic analysis revealed that ceramides accumulate in both apoptosis and senescence, whereas sphingomyelins are significantly depleted only in senescence. Gene expression data indicated an upregulation of Golgi-localized sphingomyelin synthase 1 (SMS1); however, enzymatic assays showed no significant change in its activity. To investigate the regulation of sphingomyelin levels, we inactivated ceramide transport from the ER to the Golgi via CERT, a key step in sphingomyelin synthesis. Retention of ceramides at the ER lead to a depletion of sphingomyelins and an enhanced senescence phenotype. Raman spectroscopy through the application of Ramanomics demonstrated that ceramide accumulates in the ER during senescence, supporting impaired transport to Golgi. Importantly, reducing Golgi transport of ceramides increased ER stress, suggesting that impaired ceramide trafficking contributes to the induction or maintenance of ER stress as part of the senescence program. Collectively, our findings suggest that ceramide transport from the ER is affected during senescence, contributing to ER stress activation, and supporting the role of the ER as a key site of ceramide accumulation in senescent cells.

**Scheme 1.**
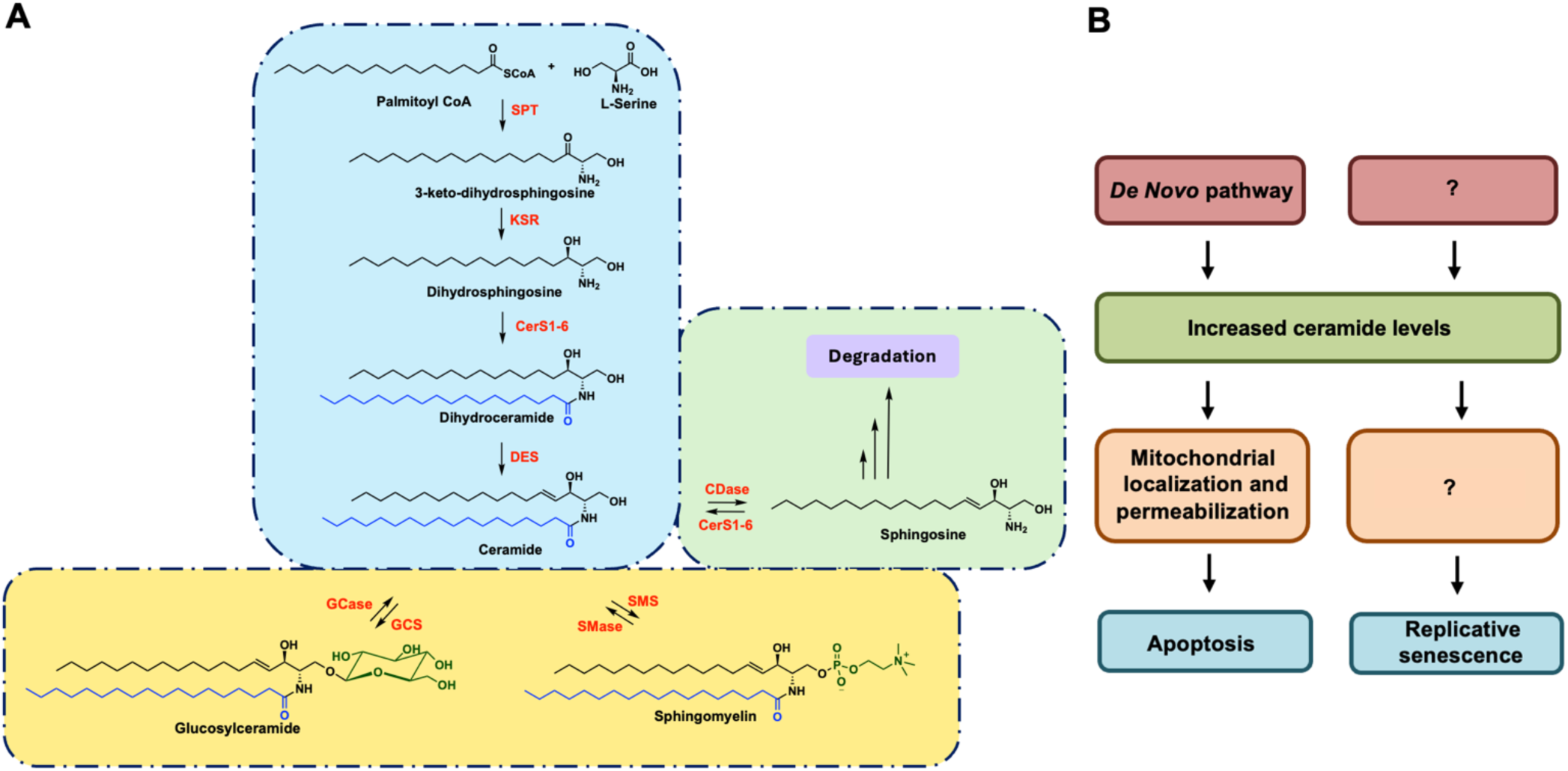
**(A)** Sphingolipid biosynthesis pathway. Simplified schematic of sphingolipid biosynthesis. Representative C18 fatty acyl chain-containing sphingolipids are shown. Enzymes involved in each biosynthetic step are indicated in red. The de novo ceramide synthesis pathway is highlighted in blue. Ceramides generated via this pathway can serve as precursors for sphingomyelin and glucosylceramide synthesis (shown in yellow) or can be further metabolized to sphingosine (shown in green). Enzymes: SPT, serine palmitoyltransferase; KSR, 3-keto-dihydrosphingosine reductase; CerS, ceramide synthase; DES, desaturase; SMS, sphingomyelin synthase; SMase, sphingomyelinase; GCS, glucosylceramide synthase; GCase, glucosylceramidase; CDase, ceramidase. **(B)** Distinct ceramide localization during apoptosis and senescence. In apoptosis, ceramides generated via de novo synthesis accumulate at the mitochondria, contributing to membrane destabilization and cell death. In contrast, ceramides also accumulate during replicative senescence; however, our findings reveal that in senescence, impaired ceramide transport results in ER-localized ceramide accumulation, contributing to ER stress and senescence maintenance.

## RESULTS AND DISCUSSIONS

### Comparative analysis of replicative senescence with apoptosis

To investigate the regulation of ceramides in replicative senescence, we employed a comparative experimental framework that integrates both senescence and apoptosis as model systems. In this context, apoptosis serves as a robust reference point for identifying senescence-specific regulatory mechanisms since ceramide dynamics are well-characterized in apoptosis. By leveraging this comparison, we aimed to uncover sphingolipid species and regulatory enzymes that are differentially modulated in senescent cells. We combined lipidomics and gene expression analyses, enabling a comprehensive investigation of sphingolipid profiles and their associated gene expression changes. We envisioned that this framework would provide a powerful platform to dissect how ceramide synthesis and metabolism diverge between apoptosis and senescence, ultimately offering insight into the distinct roles of ceramides in cellular aging and stress responses.

To carry out this comparative analysis, we used the human lung fibroblast cell line MRC-5, an established model for replicative senescence due to its well-characterized growth kinetics and senescence-associated phenotypes^13, 14^. MRC-5 cells typically undergo approximately 70 population doublings (PD) before entering the senescent state. In our experiments, cells at early population doublings (∼PD 14) were designated as “young,” while cells at ∼PD 60 were considered “senescent” based on hallmark characteristics such as increased senescence-associated β-galactosidase (SA-β-gal) activity and cessation of proliferation.^14^ We observed a linear proliferation phase up to ∼PD 35–40, after which the growth rate declined significantly, consistent with previous reports (**Figure S1A** and ref^13, 14^).

Cell pellets from both young/proliferating and old/senescent populations were collected and used for downstream lipidomic and transcriptomic profiling, following established protocols^21^. To develop a comparable apoptotic model, we treated young MRC-5 cells with doxorubicin (Dox), a DNA-damaging agent known to induce apoptosis and modulate sphingolipid metabolism. Based on dose-response viability assays, 2 µM Dox, which yielded ∼60% cell viability, was selected for further experiments (**Figure S1B**). This dual-model system, replicative senescence and apoptosis, enables a comparison of ceramide and broader sphingolipid dynamics, along with gene expression changes, across two physiologically distinct cellular states. We envisioned that this framework could uncover functional distinctions in sphingolipid metabolism during senescence and programmed cell death.

### Sphingolipid Profiling Reveals Divergent Regulatory Patterns in Senescence and Apoptosis

To investigate sphingolipid dynamics in both apoptosis and replicative senescence, we analyzed lipid profiles from senescent and apoptotic MRC-5 cells compared to their respective control groups. Lipids were extracted and analyzed as we described previously.^15, 16^ The targeted lipid analysis included major sphingolipid classes: ceramides, dihydroceramides, hexosylceramides, sphingomyelins, and dihydrosphingomyelins. We first examined total levels of each class in senescent and apoptotic cells (**Figure 1A-B**). Ceramides were significantly elevated in both conditions, with hexosylceramides following a similar trend. Interestingly, sphingomyelins exhibited opposite patterns, showing marked depletion in senescent cells (**Figure 1A**), while significantly increasing in apoptotic cells (**Figure 1B**).

**Figure 1.**
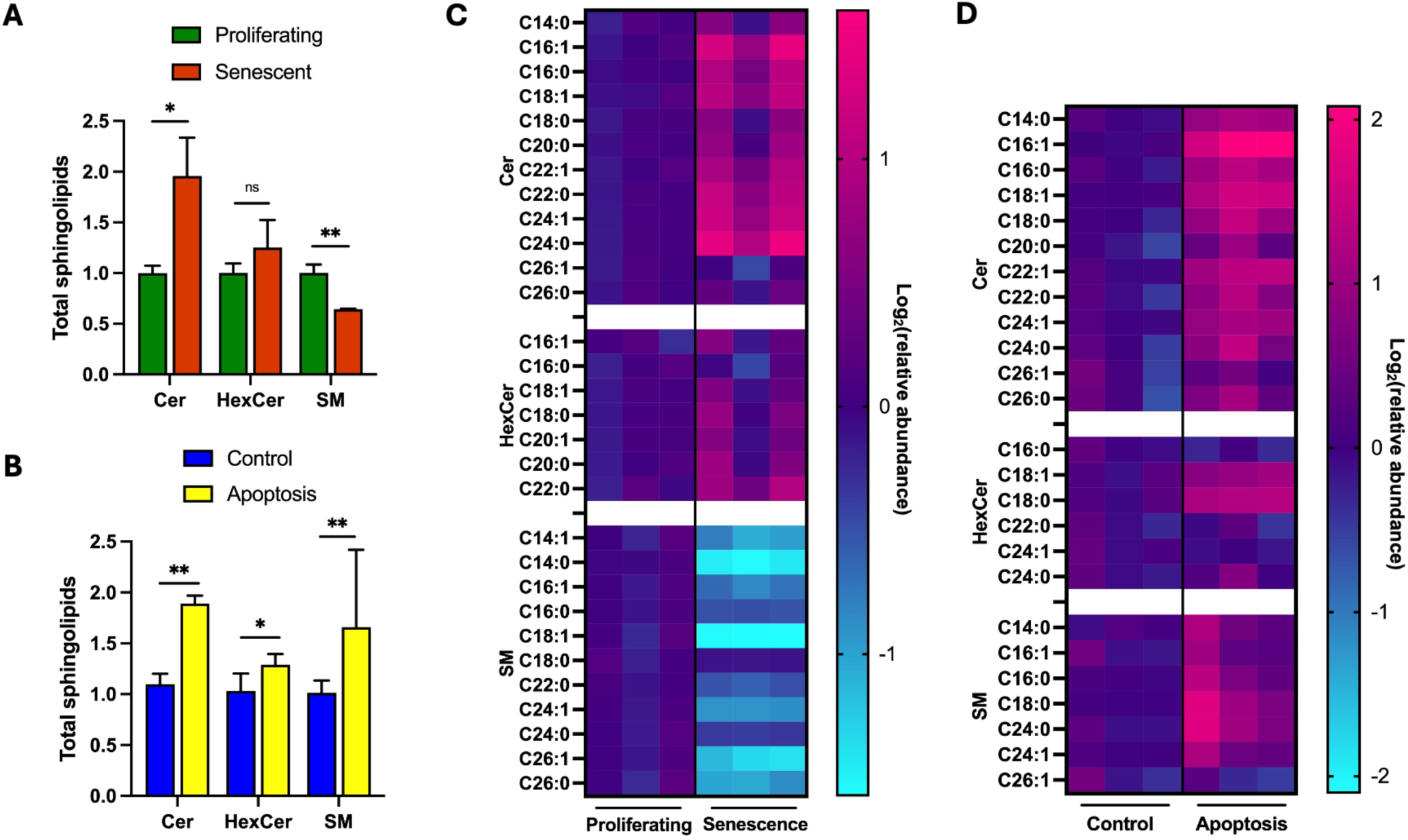
**(A-B)** Total sphingolipids levels in proliferating vs senescent and control vs apoptosis in MRC5 cells (n=3 for each condition). **(C-D)** Heatmaps representing sphingolipid levels in senescence and apoptosis model. Pink represents accumulation, and blue represents depletion. The relative abundances are reported as log_2_(relative abundances). N=3 for each condition. * represents p<0.05, ** represents p<0.01, ns represents not significant. Cer: ceramide; HexCer: hexosylceramide; SM: sphingomyelin.

To gain finer resolution, we next assessed individual sphingolipid species. Heatmaps showing relative abundances (**Figure 1C-D**) revealed distinct regulatory trends. In senescent cells, most ceramide and hexosylceramide species were significantly enriched, particularly those with long-chain (C16–C18) and very-long-chain (C20–C24) acyl groups (**Figure 1C**). In contrast, nearly all sphingomyelin species were depleted. Apoptotic cells showed elevation of ceramides and sphingomyelins, but only a subset of hexosylceramides (C18:1 and C18:0) were increased (**Figure 1D**). The divergent sphingomyelin trends are particularly striking. In senescence, the consistent depletion of sphingomyelins suggests changes in ceramide metabolism. This may be caused by perturbation of ceramide-to-sphingomyelin conversion either via changes in enzyme activity or ceramide transport.

### Gene Expression Analysis Reveals Differential Regulation of Sphingolipid Biosynthetic Enzymes in Senescence and Apoptosis

Lipidomic profiling revealed distinct sphingolipid signatures in senescence and apoptosis: ceramides were elevated in both models, but sphingomyelins showed differential regulation, significantly depleted in senescent cells while elevated in apoptotic cells. To better understand the regulation of these trends, we examined the expression of key genes involved in sphingolipid metabolism. Total RNA was extracted from MRC-5 cells representing each model (proliferating and senescent cells for the senescence model, untreated and doxorubicin-treated cells for the apoptosis model), and gene expression was quantified. We measured the expression of genes regulating *de novo* ceramide synthesis, catabolism, and downstream metabolic branching (**Figure S2**).

We then focused on genes that show differential regulation in senescence as compared to the apoptosis model. The changes in the expression trend of these genes are listed in Figure 2A. In senescent cells, the mRNA levels encoding several key enzymes or their catalytic/regulaotry subunits were significantly upregulated: Amongst these are two ER-localized proteins which drive *de novo* ceramide biosynthesis, serine palmitoyltransferase light chain 2 (SPTLC2) and dihydroceramide desaturase (DEGS1); two proteins that regulate ceramide turnover, *N*-acylsphingosine amidohydrolase 1 (ASAH1) encoding lysosomal acid ceramidase A-CDase and sphingomyelin phosphodiesterase 1 (SMPD1), encoding lysosomal acid sphingomyelinase A-SMase; two Golgi localized proteins responsible for ceramide conversion into downstream sphingolipids, sphingomyelin synthase 1 (SMS1) and glucosylceramide synthase (GCS, Figure 2A). This transcriptional signature suggests an upregulation of both ceramide production and potential utilization pathways during senescence. In contrast, apoptotic cells exhibited a distinct expression profile. While A-SMase was upregulated, genes such as SPTLC2, DEGS1, A-CDase, and SMS1 showed no significant change, and GCS was downregulated compared to controls (Figure 2A).

**Figure 2.**
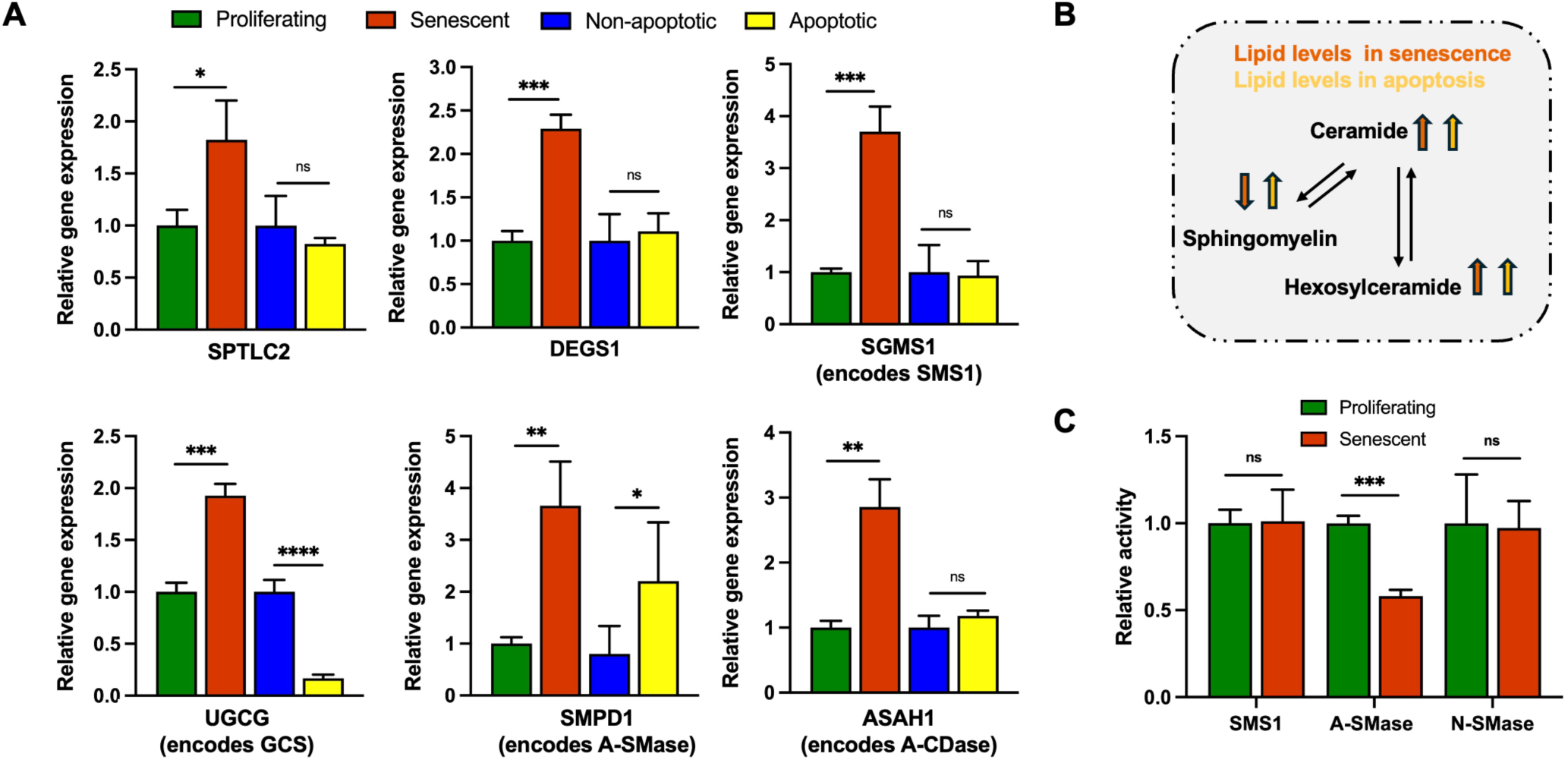
Differential regulation of sphingolipid-related genes and lipid profiles in senescence and apoptosis. **(A)** Relative gene expression levels of sphingolipid-metabolizing enzymes in senescent and apoptotic MRC-5 cells. Gene expression was normalized to the housekeeping gene HPRT1. Proteins that the genes encode for are provided in parenthesis. Data represent n = 3 biological replicates for senescent cells and n = 5 for apoptotic cells. p < 0.05 (*), p < 0.01 (**), p < 0.001 (***), p < 0.0001 (****); ns = not significant. **(B)** Schematic summary of key lipid changes observed in senescence and apoptosis based on lipidomic profiling. (**C**) SMS and N-SMase activities remain the same in senescent cells compared to proliferating ones, while a marked depletion of A-SMase is observed. The amount of C6-Sphingomyelin produced from C6-Ceramide and the amount of d_9_-C16 Cer produced from d_9_-C16 SM were measured using LC-QToF-MS.

Importantly, integration of gene expression and lipidomic data reveals a key regulatory disconnect: SMS1, the enzyme catalyzing ceramide-to-sphingomyelin conversion, is transcriptionally upregulated in senescent cells (Figure 2A-B), yet sphingomyelin levels are markedly depleted (Figure 1). Functional assays further show that SMS and N-SMase enzymatic activities remain comparable while A-SMase activity decreases in senescent cells as compared to proliferating cells (Figure 2C). These results suggest that sphingomyelin depletion may not arise from impaired enzymatic activity, but potentially from perturbed ceramide transport from the ER to the Golgi. To distinguish the involvement of enzymatic activity vs. impaired transport, we pharmacologically inhibited SMS1 using D609^22, 23^ (see Methods). D609 treatment enhanced senescence markers as measured by SA-β-gal staining (**Figure S3**). However, supplementing D609-treated cells with exogenous sphingomyelins (C2:0 sphingomyelin or C16:0 sphingomyelin, **Figure S3**) did not rescue the senescence phenotype induced by SMS1 inhibition, suggesting that it is not the reduction of sphingomyelins that cause D609-mediated senescence.

Together, these results suggest that ceramide accumulation, rather than sphingomyelin depletion, likely serves as the driver of replicative senescence in this context. These observations raise the intriguing possibility that the regulation of available ceramide for sphingomyelin synthesis, rather than ceramide synthesis per se, may be an important contributor to ceramide’s senescence-specific activity.

### Blocking Ceramide Transport from ER to Golgi Exacerbates Senescence

Our results suggest that additional factors beyond enzyme expression, such as ceramide transport, may be affecting sphingomyelin synthesis. Notably, the non-vesicular transfer of ceramides from the ER to the Golgi is mediated by CERT, which plays a critical role in ceramide trafficking.^24^ Based on these observations, we decided to target ceramide transport to the Golgi and hypothesized that a functional bottleneck at the level of CERT-mediated transport could contribute to the observed accumulation of ceramides and depletion of sphingomyelins in senescent cells.

To test this, we used HPA12, which inhibits CERT (Figure 3A).^25^ We reasoned that blocking this transport step should induce ceramide retention at the ER and phenocopy the sphingolipid changes observed in senescence. Dose-response studies identified 3 µM and 10 µM HPA12 as concentrations with minimal impact on cell viability (**Figure S4A**). Treatment of young MRC-5 cells with HPA12 led to a significant, dose-dependent increase in SA-β-galactosidase-positive cells, with 1.75-fold and 2.75-fold increases observed at 3 µM and 10 µM, respectively (Figure 3B), indicating increased senescence levels. 10 µM HPA12 treatment also reduced population doubling, supporting the senescence state (**Figure S4B**). To validate the phenotype observed with HPA12 treatment, we also employed a genetic approach by knocking down CERT using short hairpin RNA (shRNA). Two independent constructs (shCERT1 and shCERT2) reduced CERT levels by 55.3% and 73.8%, respectively (**Figure S5A)**. Consistent with the pharmacological inhibition results, CERT knockdown led to a significant increase in senescence-associated β-galactosidase activity. Specifically, shCERT1 and shCERT2 cells exhibited ∼5.3-6.7-fold higher senescence levels compared to control cells (**Figure S5B**). These findings confirm that both pharmacological and genetic disruption of CERT enhance senescence, supporting the role of CERT activity and impaired ceramide trafficking in senescence.

**Figure 3.**
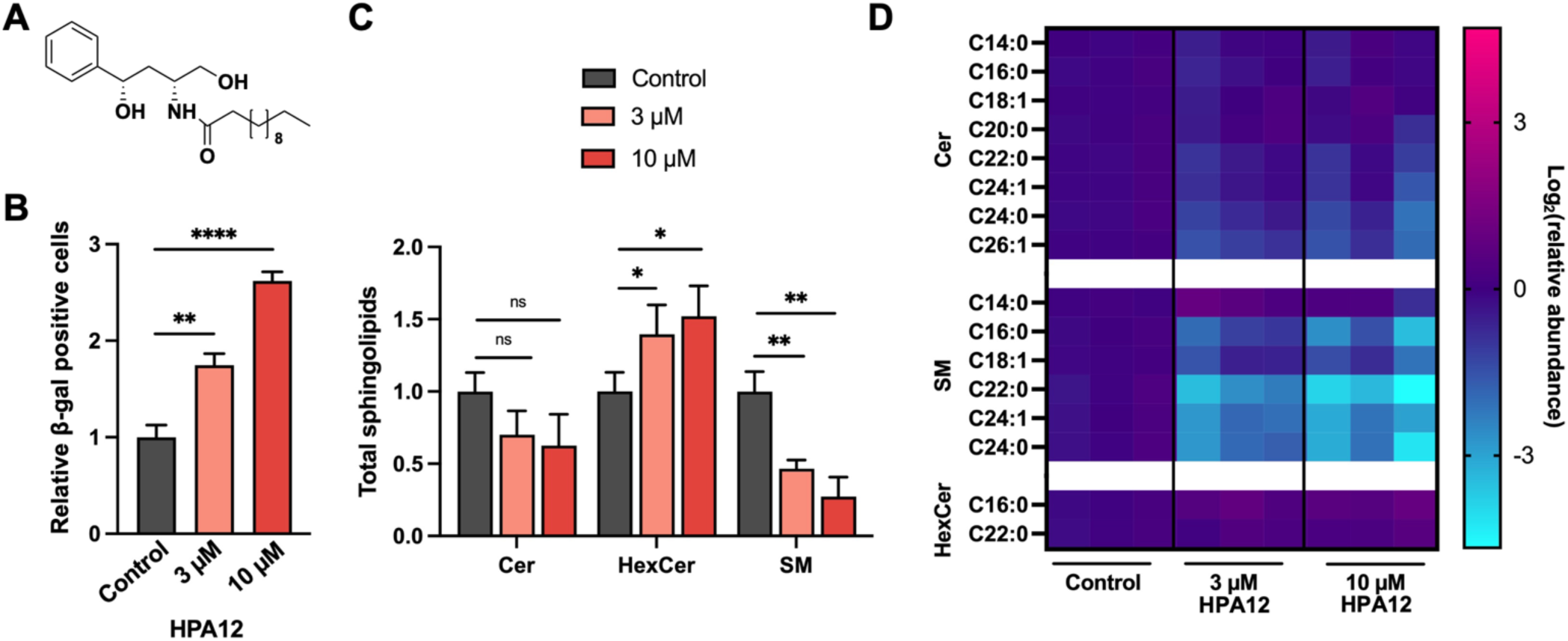
Inhibition of CERT-mediated ceramide transport and SMS enhances senescence and alters sphingolipid profiles in MRC-5 cells. **(A)** Structure of the CERT inhibitor, HPA12. **(B)** Senescence-associated β-galactosidase (SA-β-gal) assay showed an increase in senescence in MRC-5 cells treated with 3 µM and 10 µM HPA12 compared to untreated controls. **(C)** Total sphingolipid levels in MRC-5 cells following treatment with 3 µM and 10 µM HPA12, as measured by LC-QTOF-MS. **(D)** Heatmaps displaying relative abundances of individual sphingolipid species in MRC-5 cells treated with 3 µM and 10 µM HPA12. Data represent n = 3 biological replicates for senescent cells and n = 5 for apoptotic cells. p < 0.05 (*), p < 0.01 (**), p < 0.001 (***), p < 0.0001 (****); ns = not significant.

To assess the impact of CERT inhibition on sphingolipid composition, we performed lipidomic analysis on HPA12-treated cells. While total ceramide levels remained unchanged (Figure 3C), there was a clear depletion of sphingomyelins (5-10-fold, p < 0.05). The depletion of mid-, long-chain sphingomyelins is consistent with impaired CERT activity and ceramide utilization in the Golgi (Figure 3D).^26, 27^ This observation supports our gene expression data, suggesting that while the enzymatic machinery for sphingomyelin synthesis is unaffected, the transport of ceramides to the Golgi may be limiting in senescence. These findings support a model in which disrupted ceramide transport via CERT inhibition recapitulates key features of senescence, including sphingolipid imbalance and increased senescence-associated activity. They also underscore the importance of spatial control in ceramide metabolism, highlighting ceramide transport as a critical regulator of lipid remodeling during this process.

### Raman-BCA suggests increased ceramide localization at the ER during senescence

To gain insight into ceramide localization during senescence, we integrated traditional fluorescence microscopy and Ramanomics utilizing quantitative micro-Raman Spectrometry coupled to the Biomolecular Component Analysis Algorithm (Raman BCA)^28^ to monitor ceramide localization in senescent cells. Raman BCA selectively recognizes many types of biomolecules and enables studies of the molecular composition in single organelles^29, 30^. Further, in contrast to largely qualitative or semi-quantitative fluorescence measurements, the Raman-BCA approach can quantitatively determine the molecular concentrations at the site of spectra measurements in live cells.

Alkynyl probes are highly effective for Raman sensing in live cells due to their unique spectroscopic properties and biorthogonality. The carbon-carbon triple bond (-C≡C-) produces a distinct Raman signal in the cellular silent region (1800–2800 cm^−1^), minimizing spectral overlap with endogenous biomolecules and enabling high-contrast imaging. The small size of -C≡C- allows for minimal perturbation of cellular function, making them ideal for live-cell applications. The chemical versatility facilitates conjugation to a wide range of biomolecules, including proteins, lipids, and nucleic acids, allowing for targeted imaging of specific cellular structures and processes. Furthermore, unlike conventional fluorescent probes, alkynyl probes demonstrate high stability, reducing signal loss over extended imaging periods. These properties make alkynyl probes a powerful tool for studying dynamic cellular events with high specificity and minimal interference.^31^

We took advantage of the biorthogonal nature of the alkynyl group in generating the Raman signal in live cells and used C6-alkynyl ceramide and first established that alkynyl ceramide exhibits a distinctive Raman peak at ∼2100 cm^-1.30^ To minimize potential contributions from downstream metabolites, we limited treatments to one hour. In parallel experiments with C8 ceramide, we found that <10% was converted to ceramide metabolites (i.e. C8 SM or C8 hexosyl ceramide) within this timeframe, with ∼90% remaining as ceramide (**Figure S6**). Thus, the Raman signal detected under these conditions is expected to primarily reflect intact C6-alkynyl ceramide. We subsequently used C6-alkynyl ceramide signature to map the localization of ceramides in proliferating vs. senescent cells in different organelles, relevant for ceramide metabolism. We utilized LysoTracker Green DND-26, ER-Tracker Green and Mito-tracker Green to fluorescently label lysosomes, ER, and mitochondria in live cells, respectively.^32^ Data acquisition and analysis were carried out as described previously.^30^ Briefly, Raman spectra were collected from the fluorescently-labeled organelles in live proliferating and senescent cells and used to calculate the ceramide concentrations based on the peak intensity around ∼2100 cm^-1^ (Figure 4A). We found that C6-alkynyl ceramide was significantly accumulated at the ER in senescent cells compared to proliferating cells, while levels at the mitochondria were reduced, and lysosomal levels remained the same (Figure 4B). Collectively, these findings provide support for our hypothesis that ceramides are retained in the ER during cellular senescence.

**Figure 4.**
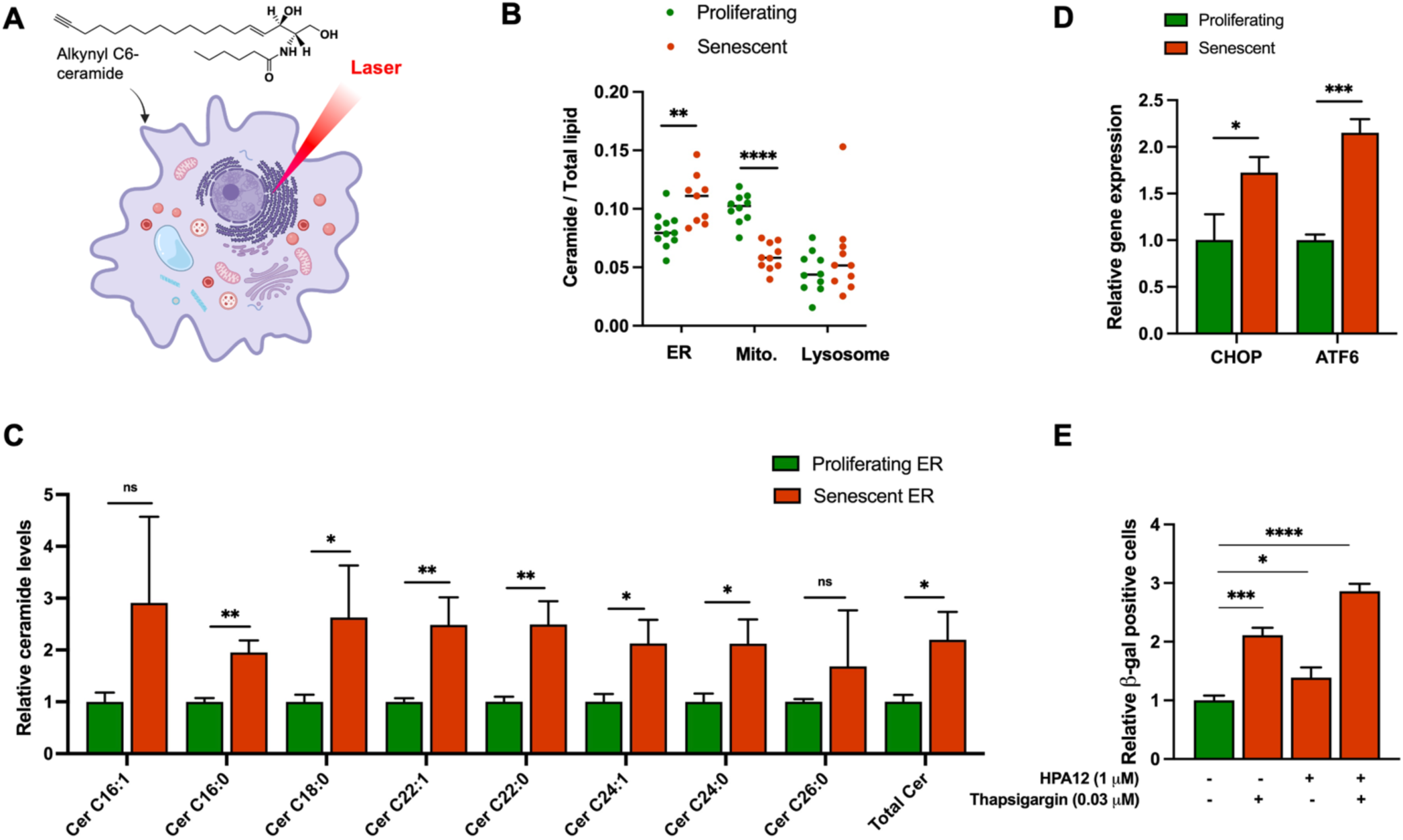
Ceramides accumulate at the ER during senescence. **(A)** Structure of alkynyl ceramide and schematic representation of Raman-BCA to understand the localization of ceramide in senescence. **(B)** Raman measurements for young and senescent cells pre-incubated with C6-ceramide alkyne probe, followed by imaging of the organelles: endoplasmic reticulum, mitochondria, and lysosome. (**C**) Bar plot showing ceramide levels in ER-enriched fractions of senescent cells relative to proliferating cells. Lipid levels are normalized to protein content of each sample (n=3 for each condition) **(D)** Relative gene expression levels for the ER stress markers: CHOP (C/EBP homologous protein), ATF6 (activating transcription factor 6) (n=3 for each condition). **(E)** Senescence-associated β-galactosidase (SA-β-gal) assay for co-treatment of HPA12 (1 µM) and thapsigargin (0.03 µM) compared to individual treatments and untreated controls (n=3 for each condition). p < 0.05 (*), p < 0.01 (**), p < 0.001 (***), p < 0.0001 (****); ns = not significant.

To further demonstrate the accumulation of ceramides at the ER during cellular senescence, we employed sucrose gradient-based ultracentrifugation to enrich for ER fractions from proliferating and senescent cells using a procedure adapted from previous work^33^ (**Figure S7**). The lipid components of these fractions were extracted using MeOH, and analyzed using LC-QToF-MS. Several ceramide species showed statistically significant accumulation in ER-enriched fractions obtained from senescent cells, overall ceramide lipids showing a two-fold increase (Figure 4C). Notably, SM levels remained unchanged in ER-enriched fractions from proliferating and senescent cells (**Figure S7B**), despite a decrease in total SM levels in senescent cells (Figure 1A), indicating that the observed increase in ER ceramides is not a consequence of a global alteration in lipid abundance but rather reflects a selective and biologically important accumulation of ceramides at the ER. These, combined with Raman-based imaging results and functional assays strongly support that ceramides accumulate at the ER due to impaired trafficking in replicative senescence.

### Ceramide-Induced ER Stress as a Potential Mechanistic Driver of Cellular Senescence

Ceramides can accumulate in the ER under metabolic and oxidative stress conditions.^34, 35^ This accumulation can perturb ER membrane composition and impair protein folding capacity, initiating ER stress and activating the unfolded protein response.^36^ Persistent ER stress has been closely associated with the onset of cellular senescence. Increasing evidence suggests a bidirectional relationship between ER stress and senescence, where ER stress not only arises as a consequence of senescence but also serves as a causal factor in its initiation and maintenance.^37^ Central mediators of the unfolded protein response, such as activating transcription factor 6 (ATF6) and C/EBP homologous protein (CHOP), have been shown to modulate senescence-associated gene expression.^38, 39^

Emerging evidence suggests that ceramide-mediated cellular stress may serve as a link between lipid metabolism dysregulation and the establishment of senescence.^40^ To determine whether ceramide accumulation at the ER contributes to ER stress during senescence, we first assessed the expression levels of key ER stress markers, CHOP and ATF6. Both markers exhibited increased expression in senescent cells compared to proliferating cells (Figure 4D), supporting increased ER stress during senescence. We then conducted a co-treatment experiment of HPA12 with thapsigargin to block the ER-to-Golgi transport of ceramide and induce ER stress, respectively^41^, and measured the senescence levels using the SA-β-gal assay (Figure 4E). Exposure to the ER stress inducer thapsigargin (0.03 µM) led to a ∼2-fold increase in SA-β-gal-positive cells compared to controls, indicating that mild disruption of ER calcium homeostasis is sufficient to elicit a robust senescence response. Pharmacological inhibition of CERT using HPA-12 yielded a significant increase in the number of senescent cells as expected based on previous results. Notably, co-treatment with thapsigargin and HPA-12 resulted in a ∼3-fold elevation in SA-β-gal positivity, surpassing the effect of either compound alone. Mechanistically, the combined effects of ER stress and ceramide accumulation within the ER may enhance unfolded protein response signaling and cellular stress and increase the proportion of cells undergoing irreversible growth arrest. Together, these findings demonstrate that both ER stress and lipid homeostatic disruption promote senescence, and their convergence amplifies the senescent phenotype, underscoring the interplay between ER stress and ceramide accumulation at the ER in senescence regulation.

## CONCLUSIONS

Ceramides are bioactive sphingolipids known to regulate diverse cellular processes, including apoptosis and senescence. While it has been established that ceramides accumulate during the senescence, the regulatory mechanisms and subcellular localization of ceramides in this process remain incompletely understood. In this study, we investigated how ceramides are distinctly regulated in replicative senescence using a comparative model of apoptosis and senescence. Through integrated lipidomics profiling, we observed that while both senescent and apoptotic cells exhibited elevated ceramide and hexosylceramide levels, sphingomyelins were significantly depleted in senescent cells but increased during apoptosis. Notably, in senescent cells, long-chain and very-long-chain sphingomyelin species were most strongly reduced. Despite increased transcription of SMS1, enzymatic activity remained unchanged, suggesting that it is not the conversion of ceramide to sphingomyelin that causes reduced sphingomyelin levels. We hypothesized that impaired ER-to-Golgi transport can mediate the changes in ceramide and sphingomyelin dynamics we observe in senescence.

To test this hypothesis, we disrupted ceramide trafficking by inactivating CERT, the ER-to-Golgi ceramide transfer protein, pharmacologically and genetically. CERT inactivation recapitulated the sphingolipid changes seen in senescent cells and significantly enhanced senescence markers in proliferating cells, supporting a model in which impaired ceramide transport and metabolism drive senescence-associated phenotypes. Using quantitative Raman-BCA and analysis of ER-enriched membranes, we also demonstrated that ceramides accumulate at the ER in senescent cells.

Ceramide accumulation at the ER has been shown to initiate ER stress responses in various models, primarily through activation of the unfolded protein response and downstream effectors.^42, 43^ Concurrently, ER stress has emerged as a potent inducer of cellular senescence, contributing to the activation of key pathways involved in growth arrest.^44^ Despite these parallel observations, a mechanistic connection linking ceramide accumulation at the ER to senescence has remained unclear. Our findings help bridge this gap by demonstrating that ceramides accumulate at the ER during replicative senescence, where they typically promote apoptosis. This ER-localized accumulation coincides with impaired ceramide flux to the Golgi, as evidenced by sphingomyelin depletion and preserved SMS1 activity despite increased expression. Together, these data suggest that ceramides are retained within the ER, where they may act as endogenous stressors that contribute to ER dysfunction and the establishment or maintenance of the senescent phenotype. Thus, our work proposes a framework in which ER-localized ceramide accumulation serves as a molecular trigger for ER stress–driven senescence. Together, these findings highlight the importance of ceramide compartmentalization and transport in regulating cell fate and identifies ceramide transport as a critical component of replicative senescence.

## SUPPLEMENTAL INFORMATION

Supplemental information includes 7 figures and 4 tables.

## ACKNOWLEDGMENTS

We acknowledge the support from the National Science Foundation grant (MCB 2314338 to G.E.A.G.) and DP2ES032761 to Y.S. M.L. is supported by the Curci Foundation. The work at the Institute for Lasers, Photonics and Biophotonics was supported by the funds from the office of Vice President for Research and Economic Development. We would like to thank Karla I. Garcia-Gonzalez for her assistance in generating lipidomics data.

## AUTHOR CONTRIBUTIONS

Experiments were designed by S.C. and G.E.A-G. The experiments were conducted by S.C., M. L., N.G., A. P., and A. K.. S.C, N.G., conducted the lipidomics, gene expression analysis and other biochemical assays. A.P., A. K. and P.P. designed and supervised Raman experiments. Y.S. designed and supervised organelle isolation experiments and provided critical feedback in data interpretation and manuscript preparation. The manuscript was written by S.C. and G.E.A-G. The study was directed by G.E.A-G.

The authors declare no conflict of interest.

## STAR METHODS

### Materials

MRC-5 cell line (human lung fibroblasts) was purchased from the American Type Culture Collection. Eagle’s Minimum Essential Medium (EMEM), penicillin-streptomycin, and trypsin were sourced from Corning. Fetal Bovine Serum (FBS) and doxorubicin were acquired from Millipore Sigma. Dimethyl sulfoxide (DMSO) was obtained from Acros Chemicals. Senescence-associated β-galactosidase staining kit was obtained from Cell Signaling. LC-MS columns and guard cartridges were purchased from Phenomenex. Lipid standards were obtained from Avanti Polar Lipids and Cayman Chemical. All solvents were LC-MS grade and purchased from Millipore Sigma. iScript Advanced cDNA conversion kit was purchased from BioRad. E.Z.N.A Total RNA kit 1 was purchased from Omega Biotek. MTT reagent was purchased from Sigma. The senescence β-galactosidase staining kit was sourced from Cell Signaling. HPA12, D609 and C6-Ceramide alkyne were purchased from Cayman Chemicals. Chloroquine and puromycin were purchased from Sigma. ER-Tracker™ Green (BODIPY™ FL Glibenclamide), LysoTracker™ Green DND-26 and MitoTracker™ Green FM Dye were bought from Invitrogen. Primary antibodies used in Western blotting: Calreticulin (Cell Signaling Technology, #12238), Tom20 (Cell Signaling Technology, #42406S), Syntaxin 6 (Cell Signaling Technology, #2869), Na,K-ATPase alpha1 (Cell Signaling Technology, #2356),), γ-Tubulin (ABclonal, A9657), Anti-rabbit IgG, HRP-linked Antibody (Cell Signaling Technology, #7074), Anti-mouse IgG, HRP-linked Antibody (Cell Signaling Technology #7076).

## Methods

### Cell Culture for senescence

MRC-5 cells, obtained at a population doubling (PD) ∼ 14 were cultured in EMEM supplemented with 10% FBS and 1% penicillin-streptomycin. Cells were grown in a humidified incubator at 37 °C and 5% CO_2_. To achieve senescence, the cells were cultured to about PD ∼ 69 (referred to as old/senescent cells hereon) in 10 cm cell culture dishes. Cell pellets were collected at population doublings 24 (young) and 69 (old).

### Doxorubicin-induced apoptosis

1 × 10^6^ MRC-5 cells were plated in a 10 cm cell culture dishes and incubated for 16 hours. After 16 hours, the cells were treated with Doxorubicin (Dox) such that the final concentration of Dox in the plate is 2 µM and the control plates were treated with an equal volume of DMSO followed by incubation for 24 hours before collecting cell pellets. Cells were collected by removing cell media in a 15 mL falcon, washing with 2.5 mL cold 1XPBS twice, and then scraping. The cell suspension was centrifuged (300 rcf, 5 min, 4 °C). PBS was removed, and the cell pellet was gently resuspended in cold 1XPBS to remove any residual media and centrifuged again (300 rcf, 5 min, 4 °C). Cell pellets were stored at -80 °C until lipid extraction was performed.

### Lipid extraction

Lipid extraction was performed using previously reported protocols^15^. Briefly, cell pellets were resuspended in 1 mL cold 1X PBS, 50 μL of which was set aside to be used for measuring protein concentration using the Coomassie (Bradford) Protein Assay Kit as per the protocol provided by the manufacturer. The remaining cell suspension in PBS was then transferred to a homogenizer to which 2 mL of chloroform and 1 mL of methanol was added. The solution was homogenized 30 times using a glass Dounce tissue grinder. The homogenized solution was centrifuged at 500 rcf, 4°C for 10 minutes to separate aqueous and organic layers. The organic layer was carefully removed and dried under a vacuum. Samples were then resuspended in chloroform spiked with internal standard based on total protein concentration. Ceramides from ER-membrane enriched fractions were extracted as we reported previously^15, 16^ using a monophasic MeOH-based extraction.

### LC-QTOF-MS Acquisition

Liquid Chromatography coupled to quadrupole time-of-flight mass spectrometry (LC-QTOF-MS) -based lipidomics analysis was performed using an Agilent 1260 high-performance liquid chromatography instrument paired with an Agilent 6530 jet stream electrospray ionization quadrupole time-of-flight mass spectrometer. Data was collected using an *m/z* range of 50-1700. For chromatographic separation, a Gemini C18 reversed-phase column (5μm, 4.6 mm × 50 mm, Phenomenex) with a C18 reversed-phase guard cartridge was used in negative electrospray mode, and a Luna C5 reversed-phase column (5 μm, 4.6 mm × 50 mm, Phenomenex) with a C5 reversed-phase guard cartridge was used in positive electrospray mode. Mobile phase A was constituted of 95:5 water:methanol (v/v) and mobile phase B was constituted of 60:35:5 isopropanol:methanol:water (v/v) in both negative and positive mode. For negative detection, 0.1% (v/v) ammonium hydroxide was added each mobile phase. For positive detection, 0.1% (v/v) formic acid and ammonium formate were added to mobile phases. The flow rate was set to 0.1 mL/min for the first 5 minutes followed by an increase to 0.5 mL/min for the remainder of the gradient. The gradient started at 5 minutes with 0% B to 100% B over 60 minutes. A shorter, 40-minute gradient was used to analyze ER extracts. Starting at 65 minutes (45 minutes for ER extracts) an isocratic gradient was applied at 100% B for 7 minutes. Equilibration of the column was followed with 0% B for 8 minutes. An electrospray ionization source was used with a capillary voltage of 3500 V and a fragmentor voltage of 175 V.

### Data Analysis for Targeted Lipidomics

For targeted lipidomics, three biological replicates of each condition in both models, young and senescent in the senescence model, control and dox-treated in the apoptosis model were used in positive and negative electron spray mode. Targeted data analysis was performed using the Agilent MassHunter Qualitative Analysis software by importing the raw data obtained from LC-QTOf-MS. The *m/z* for each ion was extracted and the peak area for each ion was manually integrated. Fold change was calculated as (Abundance _senescent_) / (Abundance _Young_) or (Abundance _Dox-treated_) / (Abundance _Control_) for each species.

### RNA extraction and cDNA Conversion

Approximately 1 × 10^6^ cells were collected for both models as described above. Total RNA was extracted for each condition using E.Z.N.A. Total RNA Kit I. The protocol provided by the manufacturer was followed. RNA concentration was measured using a Nanodrop-1000 spectrophotometer (Thermo). Three biological replicates were taken in each case. The extracted RNA was converted into its complementary DNA using reverse transcription using the iScript cDNA Synthesis kit following the manufacturer’s instructions.

### Droplet Digital Polymerase Chain Reaction (ddPCR)

Reaction mixtures were prepared in a 96-well plate by combining the cDNA, ddPCR Supermix, primers, and probes according to Bio-Rad’s protocol for the Droplet Digital PCR: QX200 System. BioRad Automated Droplet Generator was used to generate water-in-oil emulsion droplets and transferred to a new 96-well plate, which was heat-sealed using foil sheets. Target genes and reference gene, Hypoxanthine phosphoribosyltransferase 1 were amplified by thermal cycling the droplet emulsions as follows: 95 °C for 10 minutes (DNA polymerase activation), 40 cycles of 94 °C for 30 seconds (denaturation), 56 °C for 60 seconds (annealing and extension) with a final 10-minute inactivation step at 98 °C. The fluorescence of each thermally cycled droplet was measured using the QX200 droplet reader. Data was analyzed using Bio-Rad’s QuantaSoft software after threshold setting on the fluorescence of negative controls. Refer to SI Table 3 for the list of primers used to measure mRNA expression.

### SMS and SMase enzymatic activity assays

The activity assays were adapted from previous protocols.^45^ Briefly, both proliferating and senescent cells were collected, followed by lysis using probe sonication (10s on, 10s off, for 3 times) in 200 μL buffer supplemented with EDTA-free protease inhibitor (Thermo Fischer) and phosphatase inhibitor cocktail 2 (Thermo Fischer). Neutral buffer (25 mM Tris, 12 mM MgCl_2_, pH 7.5) containing EDTA-free protease inhibitor and phosphatase inhibitor cocktail 2 was used for the SMS and N-SMase enzyme assay. Acidic buffer used for A-SMase contained EDTA-free protease inhibitor and phosphatase inhibitor cocktail 2, 100 mM NaAc, 0.1% Triton X-100, pH 5. The assays were performed at least in triplicate in a 1.5 mL microcentrifuge tube, and each reaction mixture contained 80 μg of protein in 60 μL of buffer mixed with lipid standards (C6 ceramide and DOPC for SMS activity assay, and d_9_-C16 SM for N- and A-SMase). Each reaction mixture was incubated at 37 ℃ for 3 hours. The reactions were quenched using methanol. Samples were centrifuged, dried under vacuum, and resuspended in methanol. These samples were analyzed using LC-QTOF-MS targeting the lipid product.

### Sphingomyelin addback experiments

MRC5 cells were plated in 24-well plates to perform SA-β-gal assay. Cells pre-treated with 3 µM D609 (were subjected to sphingomyelin addback (using 10 µM C2 SM and 10 µM C16 SM) for 6 hours and then performed the SA-β-gal assay. We note that the cellular uptake of the exogenously added sphingomyelins were validated using LC-QToF-MS under the same experimental conditions, except using isotopically labeled C16 sphingomyelin.

### Senescence-associated β-galactosidase assay for small molecules

Approximately 7 × 10^3^ MRC-5 cells per well were seeded in a 24-well. After 24 hours of seeding, cells were treated with 3 µM and 10 µM of HPA12 and incubated for 72 hours. Senescence-associated β-galactosidase activity was assessed after 72 hours of treatment using the Senescence β-galactosidase staining kit from Cell Signaling according to the manufacturer’s instructions. After staining, cells were then washed with 1xPBS and observed under a light microscope to determine the percentage of SA-β-galactosidase positive cells (n ≥ 200 cells per condition, n=3).

### Dose-response profile and lipidomics for small molecules

MTT assays were used to assess toxicity of HPA12 and D609. About 2 × 10^4^ MRC-5 cells per well were plated in a 96-well plate and incubated for 16 hours. Cells were treated with varying concentrations of HPA12: 0 µM, 1 µM, 3 µM, 10 µM, and 30 µM, followed by 72 hours of incubation. After 72 hours, the media containing the treatment was removed from the wells, and 9% MTT solution in the media was added to the wells. The cells were incubated for 2-2.5 hours to allow the conversion of MTT to formazan crystals, which were later dissolved in DMSO. The absorbance was measured using a plate reader and was used to generate a dose-response curve. To perform the lipidomics for MRC5 cells treated with HPA12, ∼1 × 10^6^ MRC-5 cells were plated per 10 cm petri dish and incubated overnight. The next day, the cells were treated with 3 µM or 10 µM of HPA12 (n=3) and incubated for 3 days. The cell pellets were collected after 3 days and analyzed by LC-QToF MS.

### Preparation of lentiviral particles and CERT knockdown

*shCERT* in the pLKO.1 vector was obtained from Millipore Sigma as bacterial glycerol stocks *shCERT1* (NM_005713; TRCN0000315401) and *shCERT2* (NM_005713; TRCN0000382287). Plasmids were extracted using Omega Biotek E.Z.N.A. FastFilter plasmid DNA mini kit according to the manufacturer’s instructions. To prepare lentiviral particals, ∼3 × 10^5^ HEK-293T cells were plated in 6-well dishes for 16 hours. 150 μL DMEM was mixed with 100 ng VSV-G (envelope plasmid), 900 ng psPax2 (packaging plasmid), and 1 μg of the transfer plasmid (shRFP, and *shCERT*). Next, 6 μL of X-treme GENE 9 was added, and the solution was mixed by pipetting, then incubated at room temperature for 20 minutes before adding to the cells. About 5 min before transfection, the cell media was changed to fresh media supplemented with 25 μM chloroquine. The transfection mixture was added dropwise to each well and the plate was gently tilted back and forth to help ensure even distribution. Cells were incubated for 2 days, then the media was collected and filtered using a 0.45 μm pore syringe filter, then aliquoted and stored at -80⁰C.

Lentiviral transfection was carried out as we described before ^46, 47^ Briefly, ∼5 × 10^5^ cells were plated in 6-well dishes and allowed to attach overnight. Lentivirus was thawed on ice. 5 min before transfection, the cell media was replaced with fresh media containing 8 μg/mL polybrene. Lentiviral particles were then added dropwise incubated for 48 h. After selection with puromycin (2 μg/mL), cells were maintained in 1 μg/mL puromycin.

### Raman measurements and analysis

MRC5 cells were plated in a glass-bottom petri dish and incubated overnight. The next day, the media (EMEM) was changed to phenol red-free media, Opti-MEM, followed by the addition of C6-ceramide alkyne probe (final concentration, 25 μM) to both plates and incubation at 37 ℃ for 1 hour. After an hour, the organelle tracker was added to the plates and incubated for 20 minutes. Then the plates were washed with 1xPBS three times. Then Raman microscope DXR2 (Thermo-Fisher Scientific, Madison, WI), equipped with a laser source unit emitting ∼60 mW at 633 nm (ROUSB-633-PLR-70-1, Ondax), a 50 μm pinhole to shape the laser beam to a 0.7×0.7×1.5 μm3 FWHM, a Plan N 100× Olympus objective lens (NA = 1.25) and fluorescence lamp (X-Cite 120 PC, Photonic Solutions), was used to measure Raman spectra of labelled organelles. Quantitative analysis of organellar spectra was performed using the *BCAbox* software (ACIS LLC, Buffalo, NY). Detailed description of the method, output (BCA) parameters, and the calibration of Raman band intensities on the concentrations of biomolecules in the sample were described in our previous publications^30^. The alkyne has a significant intensity and a wavenumber of 2200 cm^-1^. A total of 10-12 points were imaged and analyzed for the ratio of C6-ceramide alkyne to total lipid at each point. The organelles that were imaged for understanding the localization of ceramide during senescence were: endoplasmic reticulum, mitochondria, and lysosomes.

### C8-ceramide metabolic assay

∼2 × 10^6^ were plated per 10 cm petri dish and incubated overnight. The cells were treated with 25 µM of exogenous C8-ceramide and harvested at 0, 15, 30, and 60 minutes. For each time point, lipids were extracted and analyzed using LC-QToF MS to quantify C8 ceramide-derived metabolites as we described above.

### Endoplasmic reticulum isolation from MRC5 cells

Protocol was adapted from the Basic Protocol 3 described in ref.^33^ Approximately 2 × 10^7^ cells were collected by trypsinization, washed with PBS, and resuspended in 1.5 mL HM buffer (Homogenization Medium; 0.25 M sucrose, 10 mM Tris-HCl, 1 mM PMSF, pH7.4). Cells were homogenized by 15 strokes with a tight-fitting Dounce pestle (0.025 to 0.075 mm clearance). To adjust the homogenate to 1.4 M sucrose, 3 mL of 2.0 M sucrose was added and mixed gently. The homogenate was transferred to a 14 × 89 mm UltraClear centrifuge tube preloaded with 1 mL of 1.6 M sucrose at the bottom. The sample was overlaid sequentially with 3 mL of 1.2 M sucrose and 3 mL of 0.8 M sucrose on the top. Gradients were ultracentrifuged for 2 h at 35,000 rpm in an SW41 Ti rotor using the lowest acceleration/deceleration settings. The ER-enriched band at the 1.4 M/1.6 M sucrose interface was collected, diluted with ∼4 mL HM in a 13 × 51 mm UltraClear tube, and centrifuged again for 30 min at 35,000 rpm in an SW55 Ti rotor to pellet ER membranes. The supernatant was removed, and the ER pellet was stored at −80 °C until analysis.

### Immunoblotting assay

Total cell samples and ER fractions were lysed in RIPA buffer (50 mM HEPES--KOH, 150 mM NaCl, 1% NP-40, 10 mM sodium pyrophosphate, 10 mM β-glycerophosphate, 1% sodium deoxycholate, 0.1% SDS, 2 mM EDTA, 50 mM sodium fluoride) supplemented with protease inhibitor cocktail on ice. Protein concentrations were determined using the Pierce BCA assay, and lysates were normalized accordingly. Equal amounts of protein were loaded on Bio-Rad Mini-PROTEAN TGX gels and transferred to 0.2 μm PVDF membranes using the Bio-Rad Turbo Transfer System. Membranes were blocked with 1% BSA and incubated with primary antibodies overnight at 4°C. Following by 3 times × 5 minutes washes with TBST buffer, secondary antibody incubation at room temperature for 1h and another 3 times × 5 minutes washes with TBST, the membranes were developed using Bio-Rad ECL substrate and imaged with an iBright CL1000 imager.

### HPA12 and Thapsigargin co-treatment and SA-β-gal assay

∼6 × 10^3^ were plated per well in a 24-well plate and incubated overnight. The cells pre-treated with HPA12 for 48 hrs were then co-treated with thapsigargin. We compared a total of four conditions: **(1)** vehicle treated cells (served as control), **(2)** only HPA12 (1 µM) for 72 hrs, **(3)** only thapsigargin (0.03 µM) for 24 hrs, and **(4)** co-treatment of HPA12 (1 µM) and thapsigargin (0.03 µM). After the completion of treatment, SA-β-gal assays were performed.

