## Supplemental data for "ER-Localized Ceramide Accumulation Contributes to Replicative Senescence"

**Figure S1. (A)** This is a representative growth curve for MRC-5 showing the number of days the cells were in culture along with the population doubling that was calculated each time cells were split. The population doubling is calculated using the following formula: $2^n = (P/P_0)$, where $P_0$ is the number of cells plated, P is the number of cells in the plate before splitting, n is the number of times the cell population increased by two-fold from the time they were plated to the time they were ready to be split. The x-axis at each point on the graph corresponds to the $n^{th}$ day the cells were split. The y-axis represents the population doubling of the cells at each time cells were split. **(B)** Dose-viability data for Dox. MRC-5 cells were plated in a 96-well plate and the cells were treated with varying concentrations of Dox. Cell viability was assessed using MTT assay. *** represents p<0.001, **** represents p<0.0001.

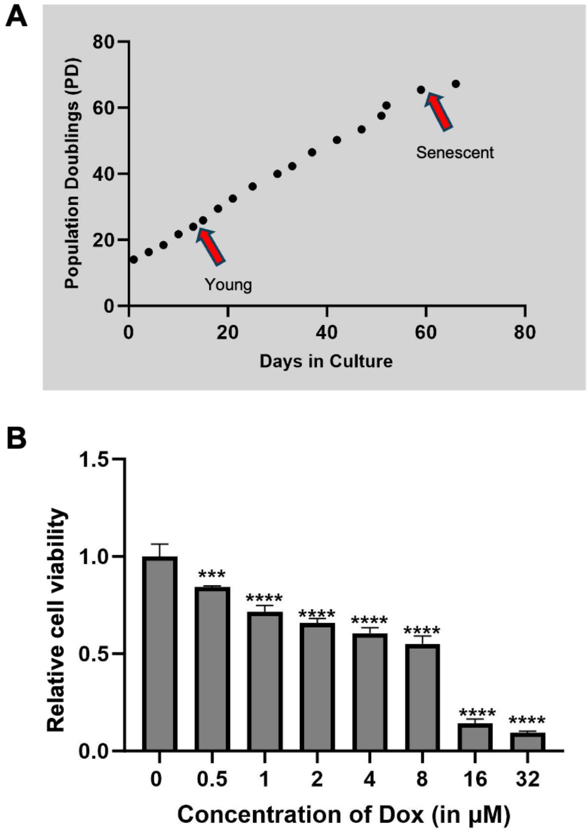

**Figure S2. Gene expression of enzymes involved in sphingolipid biosynthesis in replicative senescence and apoptosis.** This is a relative gene expression of enzymes in both models of MRC-5, human lung fibroblast cells. The expression of each gene is normalized to a housekeeping reference gene, *HPRT1*. n=3 for senescent cells and n=5 for apoptotic cells. * represents $p<0.05$, ** represents $p<0.001$, *** represents $p<0.0001$, **** represents $p<0.00001$, ns represents not significant. *SPTLC2*, serine palmitoyl transferase long chain base subunit 2; *KDSR*, 3-ketodihydrosphingosine reductase; *DEGS1*, delta 4-desaturase, sphingolipid 1; *CerS1/2/5/6*, ceramide synthase 1/2/5/6 ; *ASAH1*, N-acylsphingosine amidohydrolase 1; *SGMS1-2*, sphingomyelin synthase 1-2; *CerK*, ceramide kinase; *SMPD1-2*, sphingomyelin phosphodiesterase 1-2; *UGCG*, UDP-glucose ceramide glucosyltransferase; *GBA*, beta-glucocerebrosidase; *SPHK1-2*, sphingosine kinase 1-2; *SGPL1*, sphingosine-1-phosphate lyase 1.

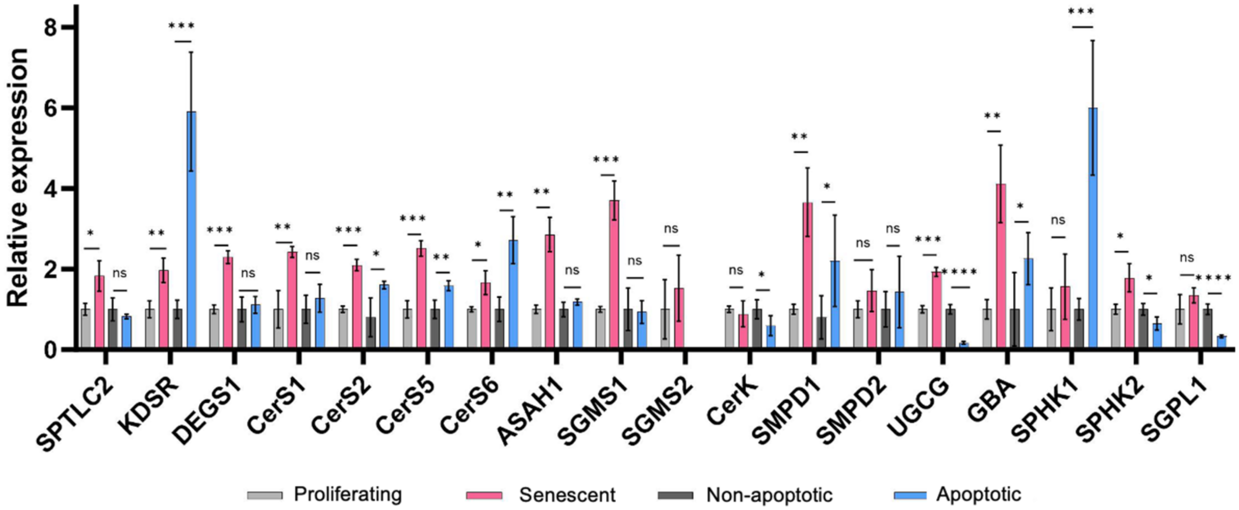

**Figure S3. Inhibition of SMS1 promotes senescence independently of sphingomyelin levels**. Pharmacological inhibition of SMS1 increases senescence markers. Exogenous sphingomyelin supplementation does not reverse the senescence phenotype, indicating that sphingomyelin depletion is not the primary driver of senescence**.** Data represent n = 3 biological replicates for senescent cells and n = 5 for apoptotic cells. $p < 0.05$ (*), $p < 0.01$ (**), $p < 0.001$ (***), $p < 0.0001$ (****); ns = not significant. Cells pre-treated with D609 were subjected to sphingomyelin addback (using C2 SM and C16 SM) for 6 hours; there was no change in the relative β-gal positive cells compared to the only D609 treatment condition.

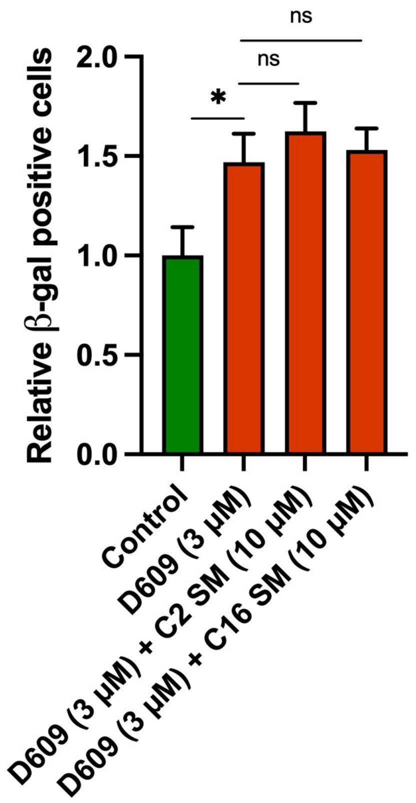

**Figure S4. Effect of HPA12 treatment on cellular viability and proliferation.** (A) Dose-cell viability data (B) Quantification of population-doubling rate following treatment. Data represent mean ± SEM from n = 3 independent experiments. **p < 0.01.

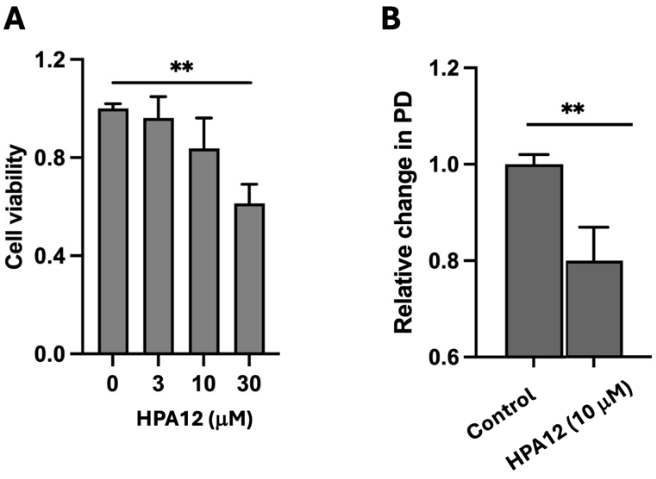

**Figure S5. *CERT* knockdown in MRC5 shows higher β-gal activity.** MRC5 cells were transfected with *shRFP*, *shCERT1,* and *shCERT2* lentiviruses and were selected with puromycin. **(A)** *shCERT1* and *shCERT2* constructs showed 56% and 74% decrease in the levels of the *CERT* gene, respectively. **(B)** shCERT1 and shCERT2 cells showed increase in β-gal positive cells, with *shCERT1* and *shCERT2* showing ~5 and ~6.7 times increase with respect to the *shRFP* cells (negative control).

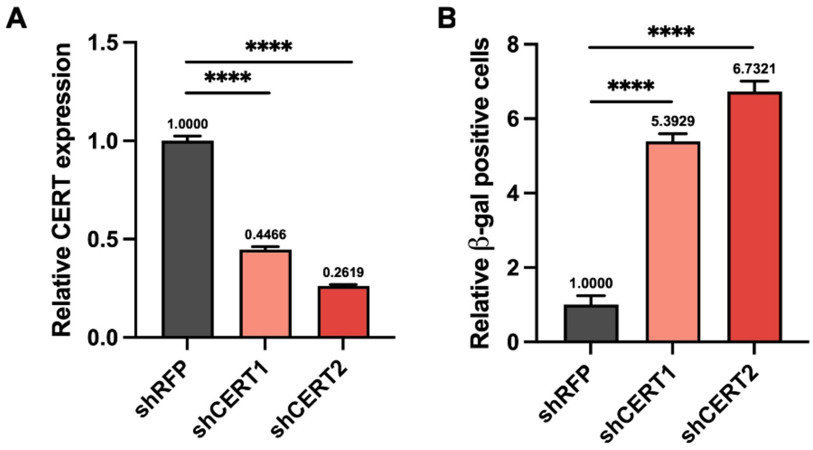

**Figure S6. C8 ceramide addition for metabolic stability.** The cells were treated with 25 μM of exogenous C8-ceramide and harvested at 0, 15, 30, and 60 minutes. For each time point, lipids were extracted and analyzed by LC-QToF MS. Percentage of ceramide remaining is reported relative to t=0 mins.

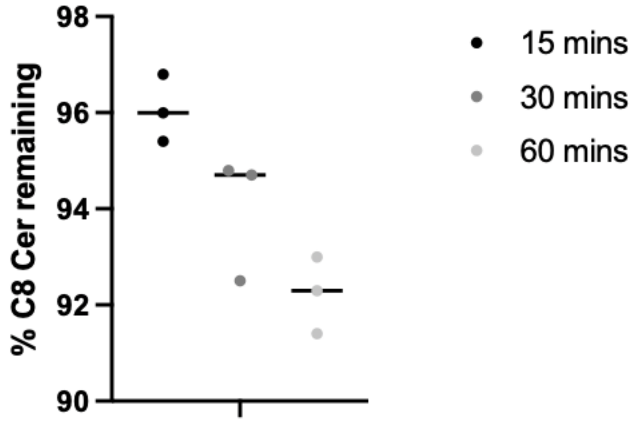

**Figure S7. Analysis of the enrichment and SM levels of fraction obtained by ultracentrifugation. (A)** Immunoblots of total MRC5 cell lysates and ER enriched lysates. Equal amounts of protein were loaded and compared. Different organelle markers were used for comparison (for ER: Calreticulin, for Golgi: Syntaxin 6, for PM: Na, K-ATPase α1, or cytosolic γ-Tubulin, for mitochondria: Tom20). **(B)** Bar plot showing SM levels in ER-enriched fractions of senescent cells relative to proliferating cells. Lipid levels are normalized to the protein content of each sample (n=3 for each condition)

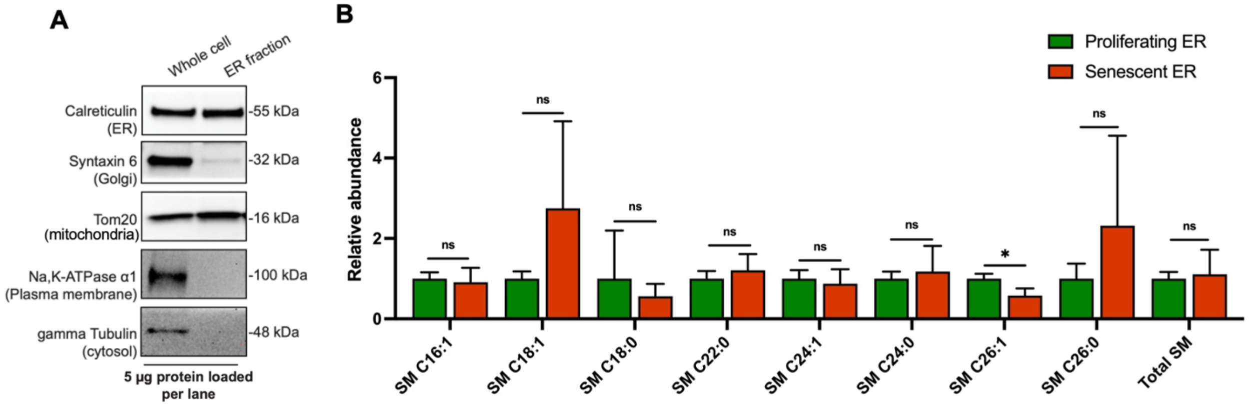

**Table S1. Sphingolipid levels in senescent cells w.r.t proliferating cells.** n=3 for each condition. m/z is the theoretical mass-to-charge ratio. Fold changes and p-values of the sphingolipid species- ceramide, dihydroceramide, hexosylceramide and sphingomyelin are provided in this table. Fold changes are calculated as **$[Abundance_{senescent\ cells}]/[Abundance_{proliferating\ cells}]$** for each species. Abundance is the total ion count for a given ion. p-value for senescent condition is calculated with respect to the proliferating cells condition.

| Species | | $m/z_{theo}$ | Ions | FC | pvalue |
|---|---|---|---|---|---|
| **Ceramide** | C14:0 | 508.4730 | $[M-H]^-$ | 1.3 | 0.1895 |
| | C16:1 | 534.4892 | | 2.3 | 0.0189 |
| | C16:0 | 536.5048 | | 1.8 | 0.0224 |
| | C18:1 | 562.5205 | | 1.9 | 0.0102 |
| | C18:0 | 564.5361 | | 1.3 | 0.1739 |
| | C20:0 | 592.5674 | | 1.5 | 0.0932 |
| | C22:1 | 618.5831 | | 1.7 | 0.0237 |
| | C22:0 | 620.5987 | | 1.9 | 0.0107 |
| | C24:1 | 646.6144 | | 2.1 | 0.0080 |
| | C24:0 | 648.6300 | | 2.5 | 0.0054 |
| | C26:1 | 674.6456 | | 0.9 | 0.5724 |
| | C26:0 | 676.6613 | | 1.1 | 0.3217 |
| **Dihydro-ceramide** | C16:0 | 538.5204 | $[M-H]^-$ | 1.2 | 0.1733 |
| | C18:0 | 566.5517 | | 1.1 | 0.3909 |
| | C20:0 | 594.5830 | | 0.9 | 0.2944 |
| | C22:0 | 622.6143 | | 1.3 | 0.0837 |
| **Hexosyl-ceramide** | C16:1 | 696.5420 | $[M-H]^-$ | 1.2 | 0.3424 |
| | C16:0 | 698.5576 | | 0.9 | 0.7287 |
| | C18:1 | 724.5733 | | 1.2 | 0.2666 |
| | C18:0 | 726.5889 | | 1.4 | 0.1453 |
| | C20:1 | 752.6045 | | 1.3 | 0.2184 |
| | C20:0 | 754.6202 | | 1.4 | 0.1735 |
| | C22:0 | 782.6515 | | 1.7 | 0.0339 |
| **Sphingomyelin** | C14:1 | 673.5279 | $[M+H]^+$ | 0.5 | 0.0064 |
| | C14:0 | 675.5436 | | 0.4 | 0.0000 |
| | C16:1 | 701.5592 | | 0.6 | 0.0029 |
| | C16:0 | 703.5749 | | 0.7 | 0.0025 |
| | C18:1 | 729.5905 | | 0.3 | 0.0016 |
| | C18:0 | 731.6062 | | 0.9 | 0.3051 |
| | C22:0 | 787.6688 | | 0.7 | 0.0017 |
| | C24:1 | 813.6844 | | 0.5 | 0.0007 |
| | C24:0 | 815.7001 | | 0.8 | 0.0313 |
| | C26:1 | 841.7157 | | 0.4 | 0.0005 |
| | C26:0 | 843.7314 | | 0.5 | 0.0079 |

**Table S2. Sphingolipid levels in apoptotic cells w.r.t control cells.** n=3 for each condition. m/z is the theoretical mass-to-charge ratio. Fold changes and p-values of the sphingolipid species- ceramide, hexosylceramide and sphingomyelin are provided in this table. Fold changes are calculated as **$[Abundance_{apoptotic\ cells}]/[Abundance_{control\ cells}]$** for each species. Abundance is the total ion count for a given ion. p-value for apoptotic condition is calculated with respect to the control.

| **Species** | | **$m/z_{theo}$** | **Ions** | **FC** | **pvalue** |
|---|---|---|---|---|---|
| **Ceramide** | C14:0 | 508.4730 | $[M-H]^-$ | 2.0 | 0.0013 |
| | C16:1 | 534.4892 | | 3.8 | 0.0020 |
| | C16:0 | 536.5048 | | 2.3 | 0.0028 |
| | C18:1 | 562.5205 | | 2.7 | 0.0013 |
| | C18:0 | 564.5361 | | 2.3 | 0.0121 |
| | C20:0 | 592.5674 | | 1.8 | 0.0599 |
| | C22:1 | 618.5831 | | 2.4 | 0.0015 |
| | C22:0 | 620.5987 | | 2.1 | 0.0221 |
| | C24:1 | 646.6144 | | 2.0 | 0.0007 |
| | C24:0 | 648.6300 | | 2.0 | 0.0590 |
| | C26:1 | 674.6456 | | 1.2 | 0.5404 |
| | C26:0 | 676.6613 | | 1.6 | 0.1255 |
| **Hexosyl-ceramide** | C16:0 | 698.5576 | [M-H]- | 0.8 | 0.2535 |
| | C18:1 | 724.5733 | | 1.8 | 0.0045 |
| | C18:0 | 726.5889 | | 2.2 | 0.0001 |
| | C22:0 | 782.6515 | | 1.0 | 0.9406 |
| | C24:1 | 808.6672 | | 0.8 | 0.2057 |
| | C24:0 | 810.6828 | | 1.2 | 0.3711 |
| **Sphingomyelin** | C14:0 | 675.5436 | $[M+H]^+$ | 1.6 | 0.1387 |
| | C16:1 | 701.5592 | | 1.4 | 0.2712 |
| | C16:0 | 703.5749 | | 1.7 | 0.1117 |
| | C18:0 | 731.6062 | | 2.3 | 0.0721 |
| | C24:0 | 815.7001 | | 2.3 | 0.0752 |
| | C24:1 | 813.6844 | | 1.6 | 0.1212 |
| | C26:1 | 841.7157 | | 0.9 | 0.6182 |

**Table S3.** Sequences of primers used for droplet digital PCR

| Gene | Description | Forward primer | Reverse primer | Exon |
|---|---|---|---|---|
| *SPTLC2* | Serine palmitoyltransferase 2 | GAAATCCAACCACAACGACAC | AATGAAGACTCTCCAGTAGTGC | 10 to 11 |
| *KDSR* | 3-Ketodihydrosphingosine Reductase | CATGGTGTCTCCGCTCATC | AGCTCCTTGTTTATAGCACTCG | 1 to 2c |
| *DEGS1* | Delta 4-Desaturase, Sphingolipid 1 | TGTAGTGAGGGAGGTTGTCAT | TTACTCATATTATGGGCCTCTGAA | 2 to 3 |
| *CerS1* | Ceramide Synthase 1 | GCCTTCCACAACCTCCTG | AACTGGGTAACAAGCAGAGTC | 6b to 6b |
| *CerS2* | Ceramide Synthase 2 | CACTGCGTTCATCTTCTACCA | GCTCTATCCTGCCTTCTTTGG | 10 to 11 |
| *CerS5* | Ceramide Synthase 5 | CCGATTATCTCCCAACTCTCAA | GCCAATTATGCCAAGTATCAGC | 8 to 9 |
| *CerS6* | Ceramide Synthase 6 | TGACTCCGTAGGTAAATACATAAAGG | CAATCAGGAGAAGCCAAGCA | 3 to 4 |
| *ASAH1* | Acid ceramidase | GCCAGTTATGACCCAGGTATC | TGTACTTCAATAGTAGCAGAAGACA | 8 to 10 |
| *SGMS1* | Sphingomyelin synthase 1 | GTACAGATAGTCCCCACACAT | GCATTTCAACTGTTCTCCGAAG | 8 to 9 |
| *SGMS2* | Sphingomyelin synthase 2 | TGCTGTACCTTCACCACTTTC | GGTCTTTCTCATCCTGCTGT | 3b to 4 |
| *CerK* | Ceramide kinase | CACCTCGCTGAACATACCATC | ACCACTGTTCACCTTAGCC | 4 to 6 |
| *SMPD1* | Sphingomyelin phosphodiesterase 1 | 5'-GAGAGAGATGAGGCGGAGA-3' | 5'-CTGGCTCTATGAAGCGATGG-3 | 2 to 3 |
| *SMPD2* | Sphingomyelin phosphodiesterase 2 | GGATCTTCAACCTCAACTGCT | GAAGTCCTGCTCACTCCA | 1 to 3 |
| *UGCG* | Ceramide glucosyltransferase | GCATTGCAACTTGAGTGGAC | GATGTGTTGGATCAAGCAGGA | 6 to 7 |
| *GBA* | Glucosylceramidase beta 1 | TTCGTTTTGCCTCCGGTT | AGAGTCTCTGAAGGAATCGAGGAT | 1 to 2c |
| *SPHK1* | Sphingosine kinase 1 | TTCACGCTGATGCTCACTG | CCGTTCACCACCTCGTG | 3 to 5 |
| *SPHK2* | Sphingosine kinase 2 | CCTTCAACCTCATCCAGACAG | CCCGTTCAGCACCTCAT | 5 to 7 |
| *SGPL1* | Sphingosine-1-phosphate lyase 1 | ATATGAGTTTGTCTTCCAGCCA | CTTGGTCTTGTTCAACTTGTCTTG | 6 to 7 |
| *CerT* | Ceramide transfer protein | CCTGAGCACGAAGATACCAAA | TGAAGATGAAACAGAGTATGGCT | 3 to 4 |
| *HPRT1* | Hypoxanthine Phosphoribosyltransferase 1 | GTATTCATTATAGTCAAGGGCATATCC | AGATGGTCAAGGTCGCAAG | 6 to 8 |

**Table S4. Sphingolipid levels in HPA12-treated cells w.r.t control cells.** n=3 for each condition. m/z is the theoretical mass-to-charge ratio. Fold changes and p-values of the sphingolipid species- ceramide, hexosylceramide and sphingomyelin are provided in this table. Fold changes are calculated as **$[Abundance_{inhibitor\text{-}treated}]/[Abundance_{control}]$** for each species. Abundance is the total ion count for a given ion. p-value for inhibitor-treated condition is calculated with respect to the control.

| **Species** | | **m/z** | **IONS** | **HPA12 (3 μM)** | | **HPA12 (10 μM)** | |
|---|---|---|---|---|---|---|---|
| | | | | **FC** | **p-value** | **FC** | **p-value** |
| **Ceramide** | C14:0 | 508.4730 | $[M-H]^-$ | 0.9 | 0.2249 | 0.9 | 0.6485 |
| | C16:0 | 536.5048 | | 0.8 | 0.2195 | 0.9 | 0.4735 |
| | C18:1 | 562.5205 | | 1.0 | 0.8817 | 1.1 | 0.5175 |
| | C20:0 | 592.5674 | | 1.0 | 0.9301 | 0.9 | 0.5840 |
| | C22:0 | 620.5987 | | 0.7 | 0.0923 | 0.6 | 0.0733 |
| | C24:1 | 646.6144 | | 0.7 | 0.0989 | 0.6 | 0.0914 |
| | C24:0 | 648.6300 | | 0.6 | 0.0281 | 0.4 | 0.0210 |
| | C26:1 | 674.6456 | | 0.4 | 0.0013 | 0.4 | 0.0036 |
| **Hexosyl-ceramide** | C16:0 | 698.5576 | $[M-H]^-$ | 1.5 | 0.0342 | 1.6 | 0.0149 |
| | C22:0 | 782.6515 | | 1.2 | 0.2200 | 1.3 | 0.0850 |
| **Sphingomyelin** | C14:0 | 675.5436 | $[M+H]^+$ | 1.6 | 0.0412 | 1.0 | 0.9372 |
| | C16:0 | 703.5749 | | 0.4 | 0.0038 | 0.2 | 0.0016 |
| | C18:1 | 729.5905 | | 0.6 | 0.0193 | 0.4 | 0.0061 |
| | C22:0 | 787.6688 | | 0.2 | 0.0044 | 0.1 | 0.0028 |
| | C24:1 | 813.6844 | | 0.2 | 0.0034 | 0.2 | 0.0026 |
| | C24:0 | 815.7001 | | 0.2 | 0.0044 | 0.1 | 0.0030 |
